# Genetic interaction between two unlinked loci underlies the loss of self-incompatibility in *Arabidopsis lyrata*

**DOI:** 10.1101/830414

**Authors:** Yan Li, Mark van Kleunen, Marc Stift

## Abstract

As the first step towards the evolution of selfing from obligate outcrossing, identifying the key mutations underlying the loss of self-incompatibility is of particular interest. However, our current knowledge is primarily based on sequence-based comparisons between selfing species and their self-incompatible relatives, which makes it hard to distinguish causal from secondary mutations. To by-pass this problem, we inferred the genetic basis of the loss of self-incompatibility by intercrossing plants from twelve geographically interspersed outcrossing and selfing populations of North-American *Arabidopsis lyrata* and determining the breeding system of 1,580 progeny. Self-incompatibility was not restored after crosses between different self-compatible populations. Equal frequencies of self-compatible and self-incompatible progeny emerged from crosses between parents with different breeding systems. We propose a two-locus genetic model for the loss of self-incompatibility in which specific *S-*locus haplotypes (*S_1_* and *S_19_*) are associated with loss of self-incompatibility through their interaction with an unlinked modifier.

## Introduction

To avoid the negative consequences of self-fertilization, about half of the angiosperms have some form of self-incompatibility^1, 2^. For example, Brassicaceae have a sporophytic self-incompatibility system, which renders plants self-incompatible through recognition and rejection of self-pollen. Two tightly linked recognition genes (the “female” gene *S*-locus Receptor Kinase – *SRK*, and the “male” gene *S*-locus Cystein Rich – *SCR*) encode stigma- and pollen proteins respectively, together forming what is commonly referred to as the *S-*locus. If stigma- and pollen proteins have matching specificities (as would be the case with self-pollination), pollen tubes cannot grow and preclude fertilization^3, 4, 5^. Self-incompatibility frequently breaks down^6, 7^, and many extant self-compatible and selfing species have evolved from self-incompatible ancestors^8, 9, 10, 11, 12^. Since the loss of self-incompatibility is the first step towards the evolution of selfing, it is of particular interest to understand its underlying genetic basis.

Given the pivotal role of self-recognition in self-incompatibility, a loss-of-function mutation at the *S*-locus could represent the primary cause of the loss of self-incompatibility. However, mutations at any of the other genes (unlinked to the *S*-locus) involved in modifying self-recognition or encoding downstream components would also render plants self-compatible^13^. Direct evidence for a primary role of the *S-*locus was obtained by a gain-of-function study that transferred a functional *S-*locus from *Arabidopsis lyrata* into *A. thaliana*, which led to complete restoration of self-incompatibility in a few accessions^14, 15, 16^. In other accessions, transformation had no effect or only led to transient self-incompatibility, indicating that these accessions must have carried loss-of-function mutations both at the *S-*locus and for additional genetic elements required for functional self-incompatibility^15^. This makes it impossible to distinguish primary from secondary mutations, because after the initial loss of self-incompatibility, relaxed selection is expected to cause pseudogenization of any locus with an exclusive function in self-incompatibility. Therefore, the finding that extant selfing species with a fully developed selfing syndrome tend to have a non-functional *S*-locus (e.g., *Capsella rubella*^17^); and *A. thaliana*^18^ does not necessarily provide evidence that mutations at the *S-*locus caused the loss of self-incompatibility. The likelihood of secondary mutations obviously increases with divergence time, which makes study systems with more recent transitions to self-compatibility more suitable to identify the mutations underlying the transitions.

Here, we make use of the intraspecific breeding- and mating system variation in North American *A. lyrata*. Since only a few *A. lyrata* populations have lost self-incompatibility and transitioned to high selfing rates^19, 20^, without evolution of a selfing syndrome^21^ or purging of genetic load^22, 23^, self-compatibility and selfing are likely of relatively recent origin. Joint clustering of selfing and outcrossing populations in multiple genetic backgrounds^20^ and an association of self-compatibility with homozygosity for two different *S*-haplotypes, *S_1_* and *S* ^24^, have provided indirect evidence for multiple origins of selfing in this species. However, direct evidence is still lacking, and it is unknown whether there is a functional link between the *S_1_* and *S_19_* haplotypes and self-compatibility. Absence of non-synonymous mutations in *SRK* and *SCR* for haplotype *S_1_* in selfing populations^25^ suggests that loss-of-function mutations at self-recognition genes cannot explain the loss of self-incompatibility. Based on segregation patterns in an F_2_ family derived from a single cross between a self-incompatible plant and a self-compatible *S_1_S_1_* plant, it was hypothesized that a recessive modifier-locus unlinked to the *S*-locus causes self-compatibility in *S_1_*-homozygotes of *A. lyrata*^24^. However, besides the single cross, there is still little empirical data to support this “modifier-hypothesis”. Moreover, it has remained unclear whether the putative modifier is specific to *S_1_*, or could also explain self-compatibility in backgrounds with other *S*-haplotypes such as *S_19_*.

To address this, we did intra- and inter-population crosses using self-incompatible plants from six predominantly outcrossing populations and self-compatible plants from six predominantly selfing populations of North-American *A. lyrata*. First, we tested whether self-incompatibility was restored in progeny resulting from crosses between self-compatible parents from different selfing populations through genetic complementation. This would be expected if two different recessive loss-of-function mutations caused the loss of self-incompatibility in different selfing populations. Second, we tested whether progeny resulting from crosses between parents with different breeding systems were always SI through functional dominance. This would be expected if a loss-of-function mutation underlies self-compatibility. Alternatively, mixtures of self-incompatible and self-compatible progeny would indicate a multigenic genetic basis of self-compatibility, for example through interaction of *S*-haplotypes and unlinked modifiers, such as proposed for the *S_1_*-haplotype^24^. Crosses within and between outcrossing populations served as controls for crosses between breeding systems, and also to test the prediction of the modifier-hypothesis that crosses between self-incompatible partners can give rise to a low frequency of self-compatible progeny, matching previous observations of a low frequency of self-compatible individuals in otherwise self-incompatible populations^24^.

## Material and methods

### Source plant material and crossing design

To study the inheritance of self-compatibility, we used seed material from 12 North American *Arabidopsis lyrata* populations (kindly provided by Barbara Mable, University of Glasgow) with contrasting breeding- and mating systems. According to genotyping of progeny arrays and self-pollinations for these 12 populations (Table S1), six populations (hereafter SI populations) mainly consist of self-incompatible (SI) plants, and have multi-locus outcrossing rates over 80%. The remaining six populations (hereafter SC populations, either *S_19_S_19_* homozygotes or *S_1_S_1_* homozygotes) have high frequencies of self-compatible (SC) plants (four populations 100%, one 88% and one 50%), and much lower outcrossing rates (five of them less than 30%, one population [TSSA] 41%; see Table S1)^20, 24^.

In March 2013, to test whether self-incompatibility could be restored through complementation, we randomly selected 18 focal plants (three plants labelled A, B and C from each of the six SC populations; Fig. 1). We used the selected parental “SC” plants to produce progeny through between-population crosses (BP_♀SC × ♂SC_) among all plants labelled A, and the same for all plants labelled B and C. As controls, we made crosses within SC populations (WP_♀SC × ♂SC_) among the A, B and C plants, respectively. To test whether self-compatibility as a trait behaves recessively, we also randomly selected 18 focal plants (three plants labelled A, B and C from each of the six SI populations; Fig. 1). We used these to produce progeny through crosses between breeding systems (BBS_♀SI × ♂SC_ and BBS_♀SC × ♂SI_) among all plants labelled A, and the same for the plants labelled B, and C. As controls for SI populations, we made crosses within SI populations (WP_♀SI × ♂SI_) among the A, B and C plants. To test whether SC plants can arise when crossing more distantly related SI plants, we also made crosses between SI populations (BP_♀SI × ♂SI_) among all plants labelled A, and the same for the plants labelled B, and C (Fig. 1). In August 2014, we replicated the complete crossing design (as summarized in Fig. 1 for the A, B, and C parental plants) with 36 different parental focal plants (three plants labelled D, E and F from each of the 12 populations).

**Fig. 1.**
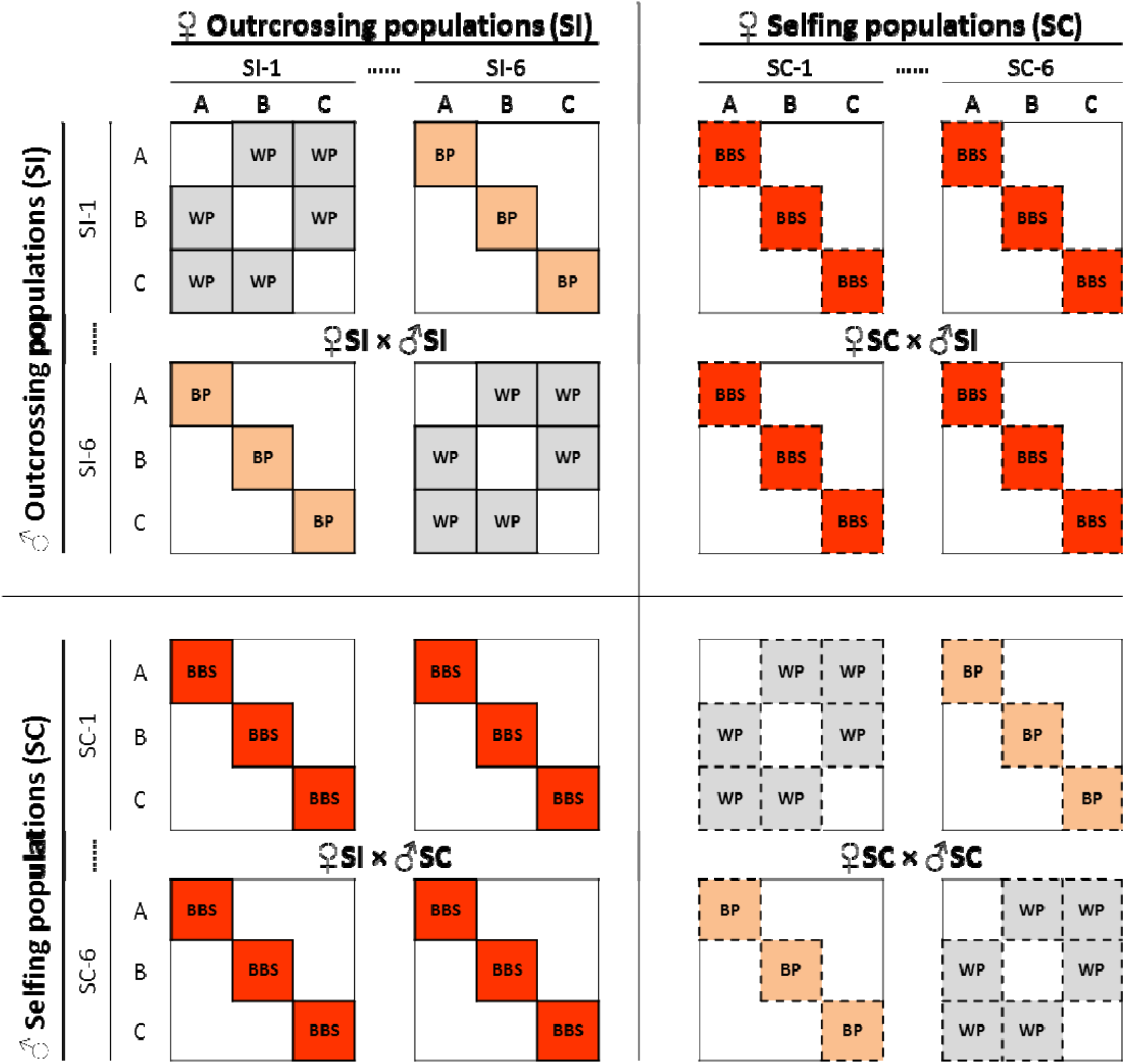
Schematic representation of the crossing design. We generated within-population (WP, grey), between-population (BP, orange; always within a breeding system) and between-breeding-system (BBS, red; always between populations) crosses using three parents (A, B, C) from six outcrossing (SI) populations (SI-1 to SI-6) and six selfing (SC) populations (SC-1 to SC-6). Solid borders indicate crosses where the maternal parent was SI, dashed borders where the maternal parent was SC. We duplicated the

In principle, our design would have generated 13 seed families per parental focal plant: 5 from crosses with plants from populations with the same breeding system (BP_♀SC × ♂SC_ or BP_♀SI × ♂SI_), 6 from crosses with plants from populations with a different breeding system (BBS_♀SI × ♂SC_ or BBS_♀SC × ♂SI_) and 2 from crosses with plants from the same population (WP_♀SC × ♂SC_ or WP_♀SI × ♂SI_). In total, for the 72 (36+36) parental plants, the complete design would thus have yielded 936 (468+468) seed families. However, not all crosses resulted in seeds, so that we obtained 898 seed families (446+452).

As not all populations are completely fixed for one or the other breeding system, we assessed the breeding system for all parental plants by performing at least six self-pollinations per plant on at least three different days. For the “SC populations”, this confirmed that all 36 plants but one (genotype D of population TSSA) were indeed SC (Table S2). For the “SI populations”, plants were either completely SI or leaky SI. Following criteria set in Refs ^19, 20^, the phenotype leaky SI is used for cases where self-pollination led to fruits with seeds, often a variable number among replicates, but fewer seeds per fruit than after cross-pollination with a compatible pollen donor.

To avoid accidental self-pollination when making crosses, we had emasculated potential recipient flowers of SC plants in the bud-stage before anther dehiscence. As a control, we left at least one emasculated flower without cross-pollen. In rare cases where this control led to fruit development (which would indicate accidental self-pollination), we discarded any fruits from flowers on the same plant cross-pollinated on the same day. To obtain at least one developed fruit for each recipient-donor combination, we did up to three cross pollinations per combination. If all three cross-pollinations did not result in fruit set, we considered the parents to be cross-incompatible. All fruits were collected when mature (4–7 weeks after pollination) and stored under dry and cool conditions until further use.

### Growth of progeny

To determine their breeding system, we sowed progeny from all 898 seed families in a growth chamber with a light period of 16 h per day, keeping temperature between 17°C and 21°C (day) and at 15°C (night) and humidity above 50%. For practical reasons, we sowed in five batches such that one seed per seed family was sown at one time, either for the 446 families derived from the A, B and C parents (batch 1, 2 and 5), or for the 452 families from the D, E and F parents (batch 3 and 4). Additionally, to increase the sample size for between-breeding-system crosses (BBS_♀SI × ♂SC_ and BBS_♀SC × ♂SI_), for this particular cross-type, we sowed seeds from all parental plants (A-F) in a sixth batch, and from parental plants A-C in a seventh and eighth batch. Batches were sown between July 2015 and February 2018 (Table S3), by placing single seeds on moistened peat-based nutrient-poor substrate (Einheitserde und Humuswerke Gebr. Patzer GmbH & Co., Sinntal, Germany) randomly assigned to a 2.5×3.2×11.0 cm cell in 54-cell QuickPot trays (QP 54 T/11, Herkuplast Kubern GmbH, Ering/Inn, Germany). Seedlings were transplanted into individual square polypropylene pots (7×7×6.5 cm, Pöppelmann, Germany) filled with the same substrate as for germination. We watered twice a week and fertilized weekly with 50 ml of 0.1% Scotts Universol® blue solution (Everris International B. V., Waardenburg, Netherlands).

### Determination of the breeding system

Although not all seeds in each batch germinated and some plants did not flower within a few months from sowing, we could perform self-pollinations on a total of 1607 progeny (Table S3) from 653 out of 898 seed families representing all 144 (12×12) population combinations. On each plant that flowered, we performed ten self-pollinations on at least five different days by rubbing a ripe anther from a donor plant on the stigma of the recipient flower.

After pollination, fruits elongate to accommodate the developing seeds, and attain their final length one to two weeks after pollination^19, 26^. Fruit length is a good proxy of seed number (Fig. S1). Therefore, as seeds can only be counted reliably at least four weeks after pollination, we used fruit length at two weeks as a proxy of seed set to enable a higher throughput and allow screening of more plants. Fruits without any developing seeds do not elongate and thus roughly maintain the length of the ovary (FL_zero_), whereas fruits with full seed set elongate to a maximum fruit length (FL_max_). Values of FL_zero_ and FL_max_ might vary among individual plants. Therefore, after pollination and fertilization, we measured the fruit lengths at least two weeks after self-pollination, and subsequently expressed the degree of self-compatibility relative to FL_zero_ and FL_max_. As it was logistically not feasible to perform control emasculations (to obtain FL_zero_ directly) and reference outcrosses (to obtain FL_max_ directly) for each progeny in the design, we used the available information from the parental plants to calculate expected FL_zero_ and FL_max_ values for each progeny. As there is considerable within and among population variation in the maximum fruit lengths, we first calculated FL_max_ for each parent individually by taking the median fruit length resulting from the four used pollen donor types (self, within population, between population, between breeding system), and taking the maximum of these medians:

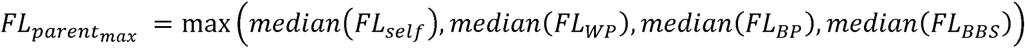

Then, given that progeny fruit length is inherited additively (Table S4), we calculated the average expected FL_max_ for each progeny

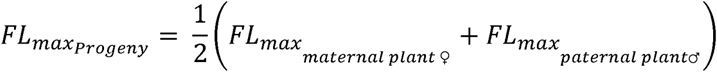

There is limited variation for the length of fruits without developing seeds within populations, but there are differences between populations^21^. To account for this, we calculated a population specific FL_zero_ as twice the mean pistil length in non-pollinated flowers for each population reported in^21^. Then, again assuming an additive contribution of the maternal and paternal parent of each progeny, we calculated the average expected FL_zero_ fruit length for each progeny:

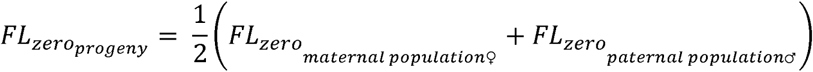

Based on these and the average fruit length resulting from self-pollination, we calculated an index of self-compatibility (SC) for each progeny:

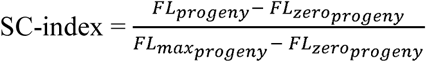

In principle, this index ranges from 0 (complete self-incompatibility) to 1 (complete self-compatibility), but is not mathematically bound by 0 and 1 due to variation in the estimates of the different component parameters.

Progeny that do not produce any elongated fruits (SC-index = 0) after self-pollination are considered to be SI, although in principle female or male sterility would give a similar outcome. To discern truly SI progeny from ones with sterility, the female and male fertility of all apparent SI progeny was tested by using them as donor (to test male fertility) and recipient (to test female fertility) in crosses with up to two haphazardly chosen unrelated progeny (only SI progeny were used if testing male fertility of the progeny). This was done for all 356 progeny that appeared SI in the first four batches. Twenty-seven out of the 356 tested progeny did not produce seeds with either partner. Although such progeny are not necessarily sterile, as cross-incompatibility could also explain lack of seed formation after crossing crosses, we conservatively excluded the 27 potentially sterile progeny from further analyses (see Table S3).

### Statistical analyses

To test for differences in SC-index between cross-types, we used linear mixed effects models implemented in the *lme* function of the ‘nlme’ package^27^ in R 3.4.3^28^. With the SC-index as the dependent variable, the model fixed part included cross-type (WP_♀SI × ♂SI_, WP_♀SC × ♂SC_, BP_♀SI × ♂SI_, BP_♀SC × ♂SC_, BBS_♀SI × ♂SC_ and BBS_♀SC × ♂SI_), and the model random part included maternal population, maternal individual (nested in maternal population), paternal population, paternal individual (nested in parental population) and batch number. To enable using a Gaussian error distribution and ensure an appropriate normality and homogeneity of model residuals, we transformed the SC-index, which ranged from −0.39 to 2.71, by adding 1.39 to all values and subsequent natural log-transformation. To account for heterogeneity of variance (i.e. differences in variance between the different cross-types), the model included a VarIdent variance structure that allowed each level of the factor cross type to have a different variance^29^.

To compare the mean SC-index among the different cross-types, we specified a matrix defining a series of 13 custom linear comparisons between different cross-type combinations (contrasts C1 to C13) and used the *glht* function in the *multcomp* package^30^ to perform z-tests corrected for multiple comparisons (Table 1). First, we tested whether progeny from between- and within-population crosses had a different SC-index, both for SI populations (C1: BP_♀SI × ♂SI_ *vs* WP_♀SI × ♂SI_) and for SC populations (C2: BP_♀SC × ♂SC_ *vs* WP_♀SC × ♂SC_). Second, we tested whether the SC-index of progeny from between-population crosses differed between SC and SI populations (C3: BP_♀SI × ♂SI_ *vs* BP_♀SC × ♂SC_). Third, we tested whether the SC-index of progeny from between-breeding-system crosses depended on the direction of the cross (C4: BBS_♀SI × ♂SC_ *vs* BBS_♀SC × ♂SI_). When C4 was not significant, we merged BBS_♀SI × ♂SC_ and BBS_♀SC × ♂SI_, and did further contrasts to test for phenotypic additivity (C5: BBS *vs* BP), complete phenotypic dominance of self-incompatibility (C6: BBS *vs* BP_♀SI × ♂SI_) or complete phenotypic dominance of self-compatibility (C7: BBS *vs* BP_♀SC × ♂SC_). When C4 was significant, we did not merge BBS_♀SI × ♂SC_ and BBS_♀SC × ♂SI_ and used separate tests for phenotypic additivity (C8: BBS_♀SI × ♂SC_ *vs* BP and C9: BBS_♀SC × ♂SI_ *vs* BP), phenotypic dominance of self-incompatibility (C10: BBS_♀SI × ♂SC_ *vs* BP_♀SI × ♂SI_ and C11: BBS_♀SC × ♂SI_ *vs* BP_♀SI × ♂SI_,), and phenotypic dominance of self-compatibility (C12: BBS_♀SI × ♂SC_ *vs* BP_♀SC × ♂SC_ and C13:BBS_♀SC × ♂SI_ *vs* BP_♀SC × ♂SC_). The SC-index values were bimodally distributed with peaks around 0 (corresponding to self-incompatibility) and 1 (corresponding to self-compatibility) (Fig. 2). Based on this, we assigned a phenotype to individual plants based on their SC-index value. Conservatively, we assigned the phenotype “SC” to all progeny with an SC-index>0.75 and “SI” to all progeny with an SC-index<0.25. Progeny with intermediate values (SC-index between 0.25 and 0.75) were thus not phenotyped, and excluded from the analyses below. To test whether self-compatibility could be explained by one or two loci, we formulated one- and two-locus models in which each locus had a single allele required for self-compatibility. Alleles required for self-compatibility could either be recessive or dominant to other allelic states, and were assumed to be fixed in SC populations (i.e., the SC cross partner was always assumed to be homozygous for the involved allele or alleles). For outcrossing populations, we used exact binomial goodness-of-fit tests to evaluate extreme scenarios in which the alleles were completely absent or completely fixed, and a “best-fit” scenario. The latter corresponds to the allele-frequency scenario that, following Hardy-Weinberg principles, best predicted the observed frequencies of SC and SI progeny from between-breeding-system crosses. For these scenarios, we also assessed the goodness-of-fit for SC and SI frequencies in progeny from crosses between SI plants within populations (WP_♀SI × ♂SI_) and between populations (BP_♀SI × ♂SI_).

**Fig. 2.**
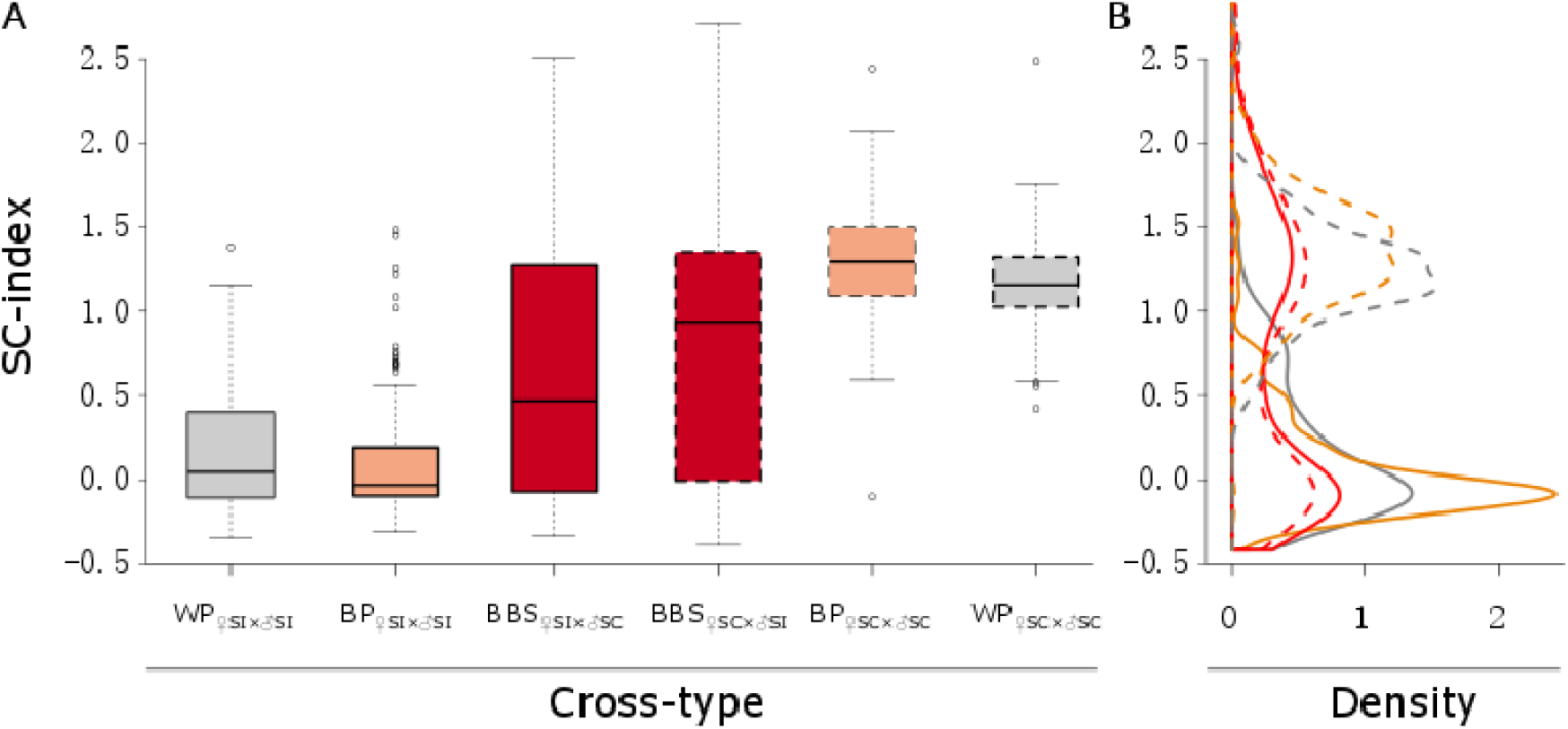
SC-index values for progeny from crosses within and between breeding systems. A) Box- and whisker plot. Boxes delimit the interquartile range (IQR) with the median indicated as a solid line. Whiskers extend to 1.5×IQR or to the lowest/highest data point within 1.5×IQR. Points beyond the whiskers’ limits represent outliers. B) Density plot. Color scheme used: grey for within-population (WP) crosses; orange for between-population (BP) crosses; red for between-breeding-system (BBS) crosses (by design also betwee populations). Solid borders and lines are used for cross-types with self-incompatible maternal parents (♀SI) and dashed borders and lines for cross-types with self-compatible maternal parents (♀SC). Note the bimodal shape for the BBS cross-type with peaks matching those observed for the cross-types from between-SI-populations crosses and between-SC-populations crosses.

**Table 1.**
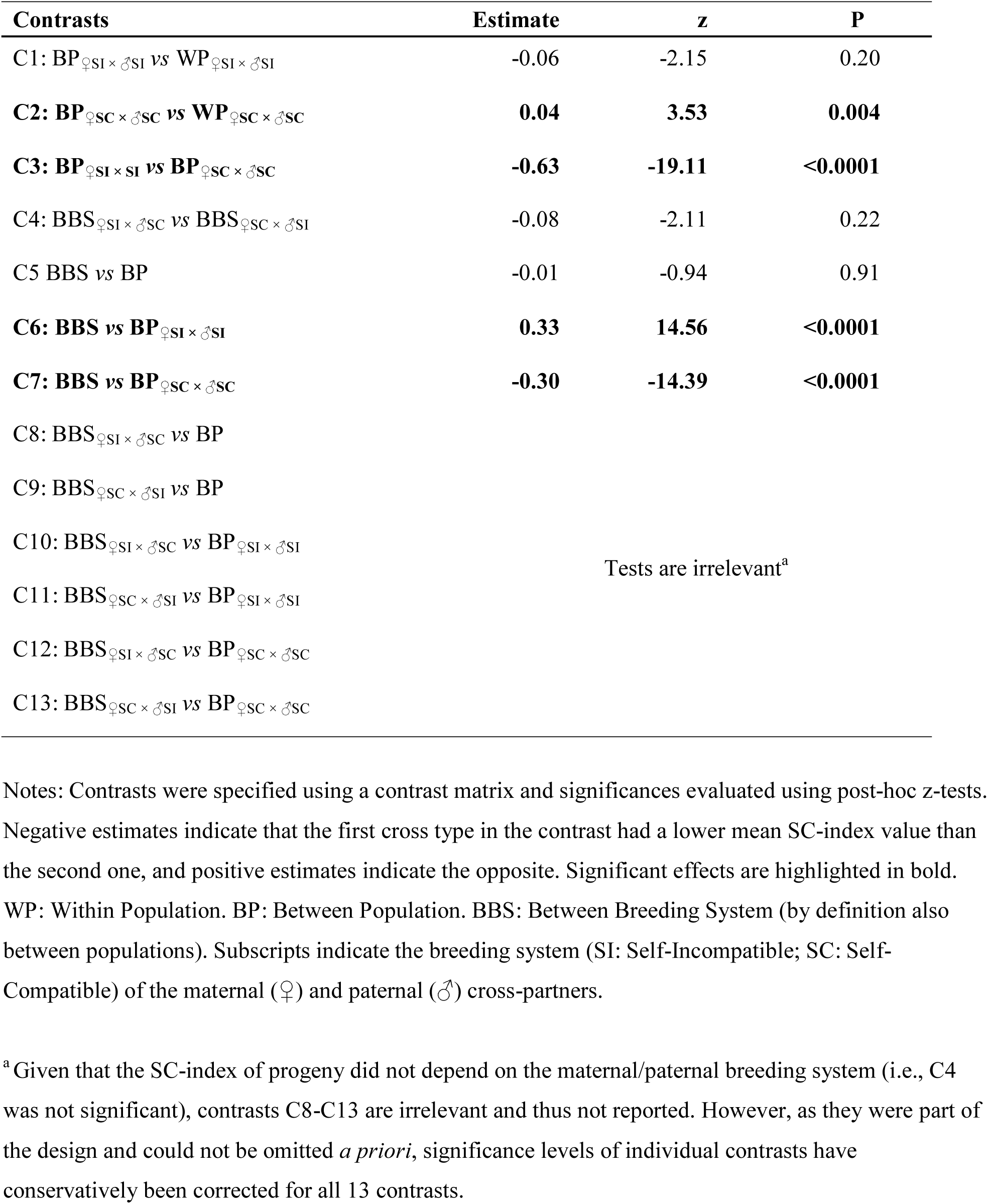
Between cross-type comparisons of mean SC-index.

## Results

### Crosses within the self-compatible breeding system (♀SC × ♂SC crosses)

For crosses within SC populations (WP_♀SC × ♂SC_), the SC-index of progeny had a unimodal distribution with a median of 1.16. The vast majority of progeny (80 out of 86) had an SC-index above the conservative threshold of 0.75, and was thus phenotyped as SC (Fig. 2, Table S5). The same pattern emerged for crosses between SC populations (BP_♀SC × ♂SC_), with an even higher SC-index median of 1.30 (significant effect of C2 in Table 1, Fig. 2), and with 233 out of 242 progeny phenotyped as SC (Table S5). Thus, overall, most progeny from crosses between SC parents showed no evidence for restoration of self-incompatibility (Fig. 3). Of the 15 cases with an SC-index below the 0.75 threshold, 14 had values above the 0.25 threshold for self-incompatibility, and could thus not be phenotyped unambiguously. Only one had an SC-index below 0.25 and was thus phenotyped as SI (Fig. 3C, Table S5). This single exception involved parents from the SC populations PTP and TSSA, which otherwise produced unambiguously SC progeny (Fig. S2).

**Fig. 3.**
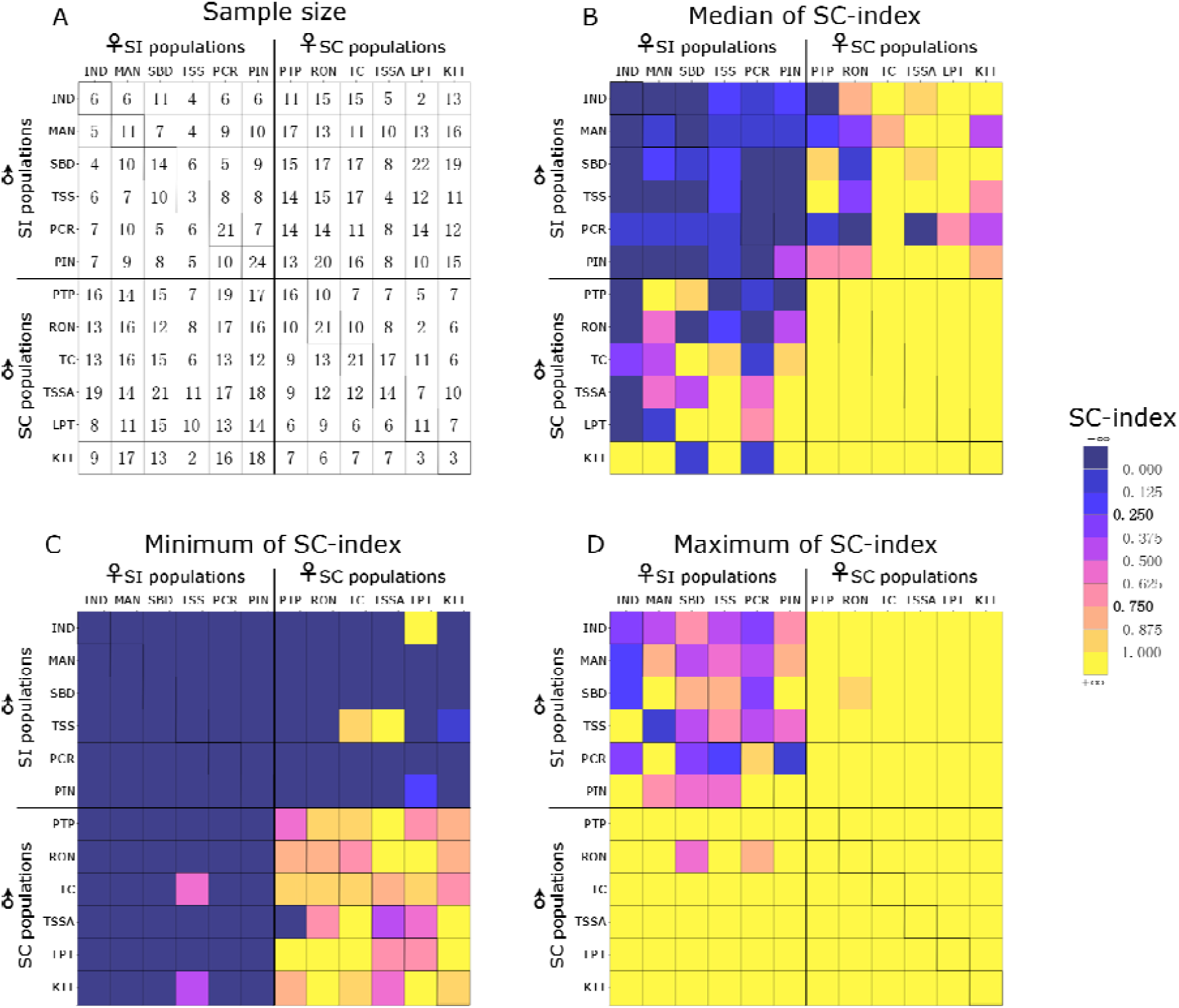
Summary statistics for the SC-index for within and between population crosses. A) Cell-specific number of progeny for which the breeding system phenotype was determined; B) Heat-map of median SC-index; C) Heat-map of minimum SC-index; D) Heat-map of maximum SC-index. Population codes correspond to the ones in Table S1. Diagonal cells marked with an extra outline represent progeny from within-population crosses. Other squares represent each possible combination of populations in two directions (with the maternal population shown at the top). The 36-cell top-left quadrat and 36-cell bottom-right quadrat represent crosses between plants with the same breeding system (♀SI × ♂SI and ♀SC × ♂SC, respectively). The 36-cell top-right quadrat and 36-cell bottom-left quadrat represent crosses between plants with different breeding systems

### Crosses within the self-incompatible breeding system (**♀**SI × **♂**SI crosses)

For crosses within SI populations (WP_♀SI × ♂SI_), the SC-index of progeny had a unimodal distribution with a median of 0.04. The majority of progeny (51 out of 79) had an SC-index below the threshold of 0.25, and was thus phenotyped as SI (Fig. 2, Table S5). However, a considerable number of progeny (9 out of 79) had an SC-index above 0.75 and was thus phenotyped as SC. The remaining 19 progeny had intermediate SC-index values and could thus not be phenotyped unambiguously according to our (conservative) thresholds (Table S5). Similar patterns emerged for crosses between SI populations (BP_♀SI × ♂SI_, no significant effect of C1 in Table 1). With a median SC-index of −0.03 (Fig. 2), 171 out of 215 progeny was phenotyped as SI, 8 as SC and the remaining 36 intermediate (Table S5). Thus, overall, most progeny from crosses between SI parents were SI (Fig. 3). However, SC progeny emerged in crosses within four out of six SI populations (MAN, PCR, PIN, SBD), and in several crosses between SI populations (eight different population-combinations; Fig. 3D).

### Crosses between plants with a different breeding system (**♀**SC × **♂**SI and **♀**SI × **♂**SC crosses)

The range of SC-index values of progeny from between-breeding-system crosses (BBS _♀SI × ♂SC_ and BBS_♀SC × ♂SI_) included the complete spectrum from completely SI to completely SC. Although on average intermediate to the SC-index of progeny from crosses within breeding systems (BP_♀SC × ♂SC_ and BP_♀SI × ♂SI_; no significant effect of C5 in Table 1), most between-breeding-system progeny could be phenotyped as either SC (477 out of 958) or SI (382 out of 958) (Table S5). Consequently, the SC-index showed a bimodal distribution with a median of 0.74 and peaks at c. 0 and c. 1.2 (Fig. 2B). The remaining 99 progeny had intermediate SC-index values and could thus not be phenotyped unambiguously according to our (conservative) thresholds (Table S5). The cross direction did not have a significant effect on the average SC-index value of progeny from between-breeding-system crosses (no significant effect of C4 in Table 1; Fig. 2), although the proportion of SC plants was higher when the mother was SC (Table S5). The patterns were not specific to any cross-combination, as the vast majority of population-combinations (31 out of 36) resulted in both SI and SC progeny (Fig. 3C; Fig. 3D). At the seed family level, 103 out of 266 families segregated for breeding system, i.e., they contained both SI progeny (SC-index<0.25) and SC progeny (SC-index>0.75) (Fig S2).

#### Evaluation of one and two-locus models to explain the loss of self-incompatibility

The observed relatively equal frequencies of SC and SI plants strongly speak against a single-locus genetic basis with completely additively interacting alleles, as this should have resulted in a unimodal distribution of SC-index values after between-breeding-system crosses. Similarly, single-locus models with allelic dominance of the allele conferring self-compatibility are unlikely, as these should have resulted in phenotypic dominance of self-compatibility, which could be statistically rejected (significant contrasts C6 and C7 in Table 1; significant deviation from single-locus dominant model in Table 2). The only single-locus model that could fit our observations would be one where the mutation acts recessively and has a relatively high frequency of 0.555 in SI populations (Table 2). However, for that scenario to work, crosses between SI parents should have yielded a much higher frequency (30.8%) of SC individuals than observed both within and between populations (15% and 4.5%, respectively; Table 2).

**Table 2.**
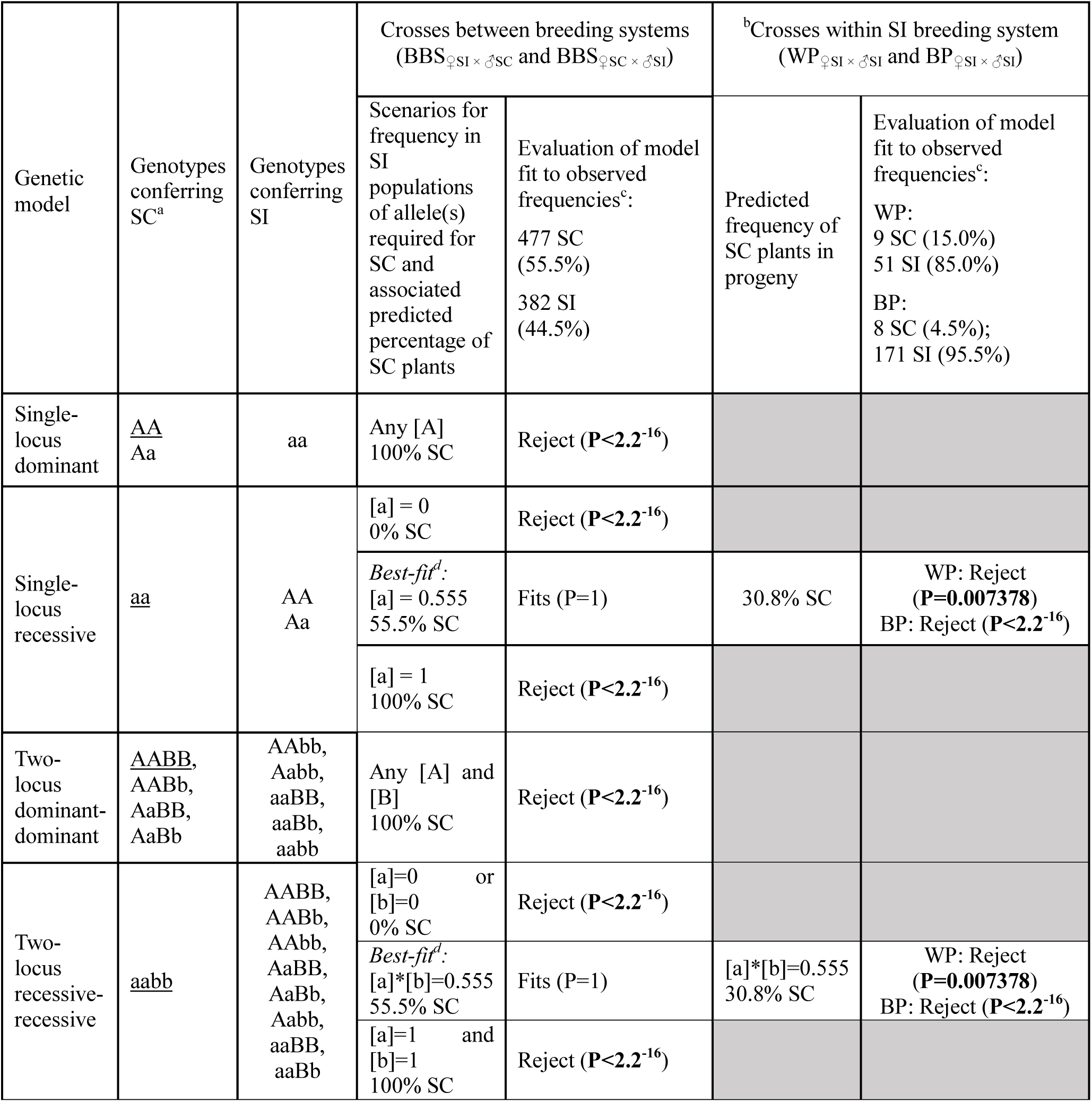
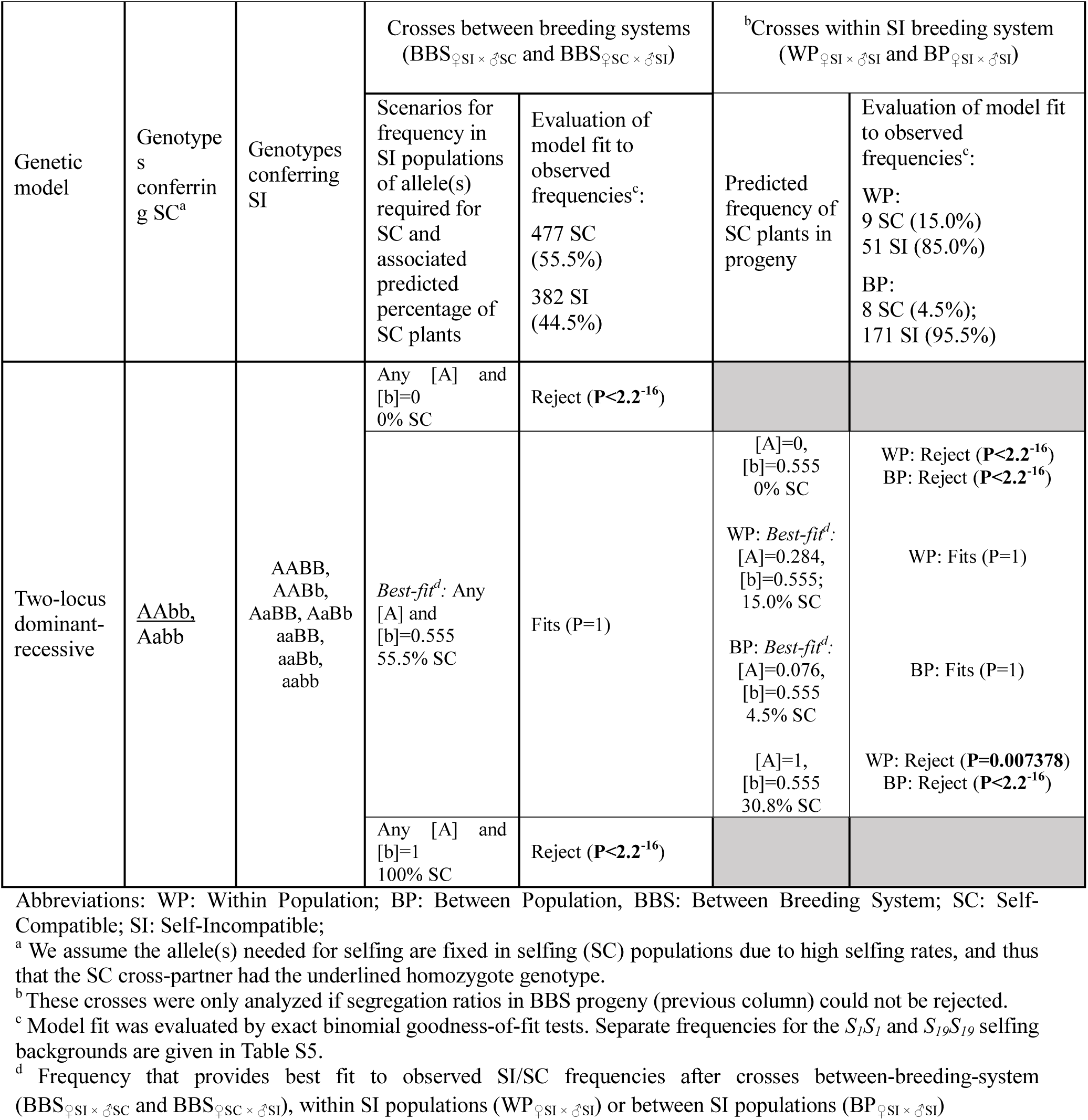
Evaluation of one- and two locus genetic models to predict breeding system segregation ratios.

Based on the overall observed frequencies of SC and SI plants after between-breeding-system crosses, two-locus models with dominance of both alleles required for self-compatibility could be rejected regardless of the assumed allele frequencies in outcrossing populations. Two-locus models in which one or both alleles act recessively could predict our data if the product of the frequency of the alleles required for SC equaled 0.555 (for the recessive-recessive models), or if the frequency of the recessive allele equaled 0.555 (for the dominant-recessive models).

Based on the observed frequencies of SC and SI plants after crosses between SI plants, the recessive-recessive model could be rejected, because the frequency of SC individuals (15% for within population crosses; 4.5% for between population crosses) did not fit the prediction of 30.8% SC progeny (Table 2). The dominant-recessive model could predict our data at the inferred recessive allele-frequency of 0.555, if the required dominant allele had a frequency of 0.284 (for crosses within populations) or 0.076 (for crosses between populations).

## Discussion

By determining the breeding system of 1,580 progeny from crosses within and between selfing and outcrossing populations of North American *Arabidopsis lyrata*, we obtained unique insights in the genetic basis of the loss of self-incompatibility in a species with a relatively recent (intraspecific) transition to selfing. Our first main finding is that most progeny from crosses between SC parents from six different selfing populations were also SC. This shows that the loss of self-incompatibility in the different selfing populations of *A. lyrata* cannot be explained by different recessive loss-of-function mutations. Our second main finding is that crosses between parents with different breeding systems yielded an even segregation ratio of breeding-system phenotypes (i.e., equal frequencies of SC and SI progeny). Taken together, our findings show that single-locus models cannot explain the genetic basis of the breakdown of self-incompatibility in North American *A. lyrata*. Instead, we propose that at least two loci are involved in self-compatibility: the *S*-locus and one or more modifier-loci. More specifically, we propose that plants carrying the dominant *S*-haplotype *S_19_* are rendered SC if they are homozygous for a recessive modifier-allele, and that plants homozygous for the recessive *S*-haplotype *S_1_* are rendered SC if they carry a dominant modifier-allele.

A single loss-of-function mutation in any of the genes required for self-incompatibility provides the simplest potential genetic mechanism that would explain the breakdown of self-incompatibility^13^. As it is improbable for the same mutation to happen twice, under a scenario of multiple origins of self-compatibility, one would then expect that different loss-of-function mutations underlie self-compatibility in different selfing populations. Indeed, different loss-of-function mutations at the *S-*locus have been found in different *A. thaliana* haplogroups^31^. Since loss-of-function mutations are usually recessive^32^, one would thus expect restoration of self-incompatibility through genetic complementation when the genomes of two SC plants with different recessive loss-of-function mutations are combined. Our results, however, do not support this scenario for *A. lyrata*. When crossing SC plants from six selfing populations with three different population genetic backgrounds^20^, self-incompatibility was never restored. This lack of genetic complementation could either imply that one and the same recessive loss-of-function mutation causes self-compatibility in the different selfing backgrounds, or that the loss of self-incompatibility has a more complex genetic basis involving more loci.

Our finding of roughly equal frequencies of SI and SC progeny from between-breeding-system crosses (BBS_♀SI × ♂SC_ and BBS_♀SC × ♂SI_) and the emergence of some SC progeny from crosses between SI plants (WP_♀SI × ♂SI_, BP_♀SI × ♂SI_) supports the involvement of at least two loci in conferring self-compatibility in *A. lyrata*. This could reflect that a genetic interaction underlies the loss of self-incompatibility, but could also be explained by secondary mutations accumulating after a single primary causal mutation. Such accumulation is expected due to relaxed purifying selection on genes involved in self-incompatibility after a transition to selfing, especially in selfing species that have diverged from their outcrossing ancestors sufficiently long ago for a selfing syndrome to evolve^33^. The resulting molecular signatures of secondary mutations in *A. thaliana*^18^ and *C. rubella*^17^ make it difficult to distinguish primary from secondary mutations in such species^13^. However, in the case of *A. lyrata*, secondary mutations are much less likely given the recent intraspecific origin of selfing in this species. Therefore, the involvement of two loci in *A. lyrata* more likely reflects a genetic interaction in which an allele of one locus confers self-compatibility by interacting with one or more alleles of another locus.

Interactions of modifiers with the *S*-locus are common features of homomorphic self-incompatibility systems^34, 35, 36^. Modifiers are particularly important in regulation of the sporophytic self-incompatibility of the Brassicaceae^13, 37, 38^. Based on this idea, it was proposed that in *A. lyrata*, an allele for a modifier-locus confers self-compatibility in plants homozygous for the *S-*locus haplotype *S_1_*^24^. However, the empirical support for this model was thin, as it was only based on segregation ratios in a single F_2_family derived from one ♀SI × ♂SC cross. Moreover, although two selfing populations are fixed for the *S_1_* haplotype, four of the selfing populations are fixed for the *S_19_* haplotype^24^. Our comprehensive crossing design, involving pairwise crosses between representatives from all six known selfing populations and six outcrossing populations, provides the first rigorous test of the modifier-hypothesis. The observed segregation of SC and SI cross-progeny in the first generation after between-breeding-system crosses, and the emergence of SC progeny after crosses between SI plants can all be explained by the hypothesis. Together, this provides strong support that self-compatibility in all selfing populations of North American of *A. lyrata* is conferred by a genetic interaction between modifier-alleles and the *S*-locus haplotypes *S_1_* and *S_19_*.

The appearance of SC progeny in crosses between SI parents from different populations suggests that one or more of the modifier-alleles segregate in outcrossing populations. Maintenance of modifier-alleles conferring partial or complete self-compatibility is expected as a means to provide reproductive assurance under conditions where costs of complete self-incompatibility are high^39, 40, 41^. As the *S_19_-*haplotype is dominant and thus relatively rare (10%) in our six SI populations^24^, the modifier allele interacting with *S_19_* should act recessively, and have a relatively high frequency in outcrossing populations (>50%) to explain the observed equal frequencies of SI and SC phenotypes in between-breeding-system crosses. Unlike the dominant *S_19_*, *S_1_* is recessive to any other *S*-haplotype^42^, and thus generally has a high frequency (79%) in outcrossing populations (e.g., Refs ^24, 43^). Thus, unless there is segregation distortion against *S_1_*^44^, this would mean that about 60% of the plants in outcrossing populations are homozygous for *S_1_*. Since SC plants are relatively rare in outcrossing populations, this implies that the modifier-allele conferring self-compatibility in *S_1_* homozygotes should have a relatively low frequency in outcrossing populations (much lower than the inferred frequency of the modifier conferring self-compatibility in plants carrying *S_19_*). Based on this, we propose that *S_1_* and *S_19_* do not interact with the same modifier-allele. This would also explain why between-breeding-system crosses yielded different frequencies of SC phenotypes depending on whether the SC parent came from a selfing population fixed for *S_1_* or from a population fixed for *S_19_* (Fig. S3; Table S5). It remains to be tested whether the two modifier-alleles are variants of the same locus, or belong to independent loci, and whether they can also interact with other *S-*locus haplotypes.

## Acknowledgements

We thank Barbara Mable for providing seeds from natural populations, Sina Konitzer-Glöckner, Katya Mamonova and Nadja Köhler for help with crossing, and Heinz Vahlenkamp, Karoline Jetter, Yanjie Liu, Samuel Fernandes and Beate Rüter for help with the experiment. YL was funded by a scholarship from the China Scholarship Council. MS was supported by grant 388824194 from the German Research Foundation (Deutsche Forschungsgemeinschaft).

## Author contributions

MS conceived the study. MS and MvK designed the experiment. YL executed the experiment. YL analyzed the data with inputs from MS and MvK. YL and MS wrote the paper with input from MvK.

## Supporting information

**Table S1.** Background information for the six selfing and six outcrossing *A. lyrata* populations from which we used seeds.

| Population code | Location, Lake, State/Province, Country | Population coordinates | | Predominant breeding system (proportion of SC individuals)<sup>a</sup> | Outcrossing rate $T_m^a$ |
| --- | --- | --- | --- | --- | --- |
|  |  | Latitude | Longitude |  |  |
| **IND** | Indiana Dunes National Lakeshore, Lake Michigan, Indiana, USA | N 41°37'17" | W 87°12'44" | SI (0.14) | 0.99 |
| **MAN** | Manitoulin Island, Georgian Bay, Ontario, Canada | N 47°39'54" | W 82°15'52" | SI (0.13) | 0.83 |
| **SBD** | Sleeping Bear Dunes National Lakeshore, Lake Michigan, Michigan, USA | N 44°56'20" | W 85°52'13" | SI (0.00) | 0.94 |
| **TSS** | Tobermory Singing Sands, BPNP, Lake Huron, Ontario, Canada | N 45°11'33" | W 81°35'02" | SI (0.00) | 0.91 |
| **PCR** | Port Crescent State Park, Lake Huron, Michigan, USA | N 44°00'15" | W 83°04'26" | SI (0.00) | 0.98 |
| **PIN** | Pinery Provincial Park, Lake Huron, Ontario, Canada | N 43°16'08" | W 81°49'53" | SI (0.00) | 0.84 |
| **PTP** | Point Pelee National Park, Lake Erie, Ontario, Canada | N 41°55'40" | W 82°30'51" | SC (1.00) | 0.09 |
| **RON** | Rondeau Provincial Park, Lake Erie, Ontario, Canada | N 42°15'41" | W 81°50'47" | SC (1.00) | 0.28 |
| **TC** | Tobermory Cliffs, Bruce Peninsula National Park, Georgian Bay, Ontario, Canada | N 45°14'30" | W 81°31'03" | SC (0.88) | 0.18 |
| **TSSA** | Tobermory Singing Sands Alvar, BPNP, Lake Huron, Ontario, Canada | N 45°11'27" | W 81°35'26" | SC (0.50) | 0.41 |
| **LPT** | Long Point Provincial Park, Lake Erie, Ontario, Canada | N 42°34'47" | W 80°23'15" | SC (1.00) | 0.13 |
| **KTT** | Kitty Todd State Nature Preserve, -, Ohio, USA | N 41°37'14" | W 83°47'15" | SC (1.00) | 0.31 |

Notes: Seeds from natural populations were kindly provided by Barbara Mable allowing us to grow the parents of our crossing design (Fig. 1).
<sup>a</sup> Values from Ref<sup>20</sup>

**Table S2.**
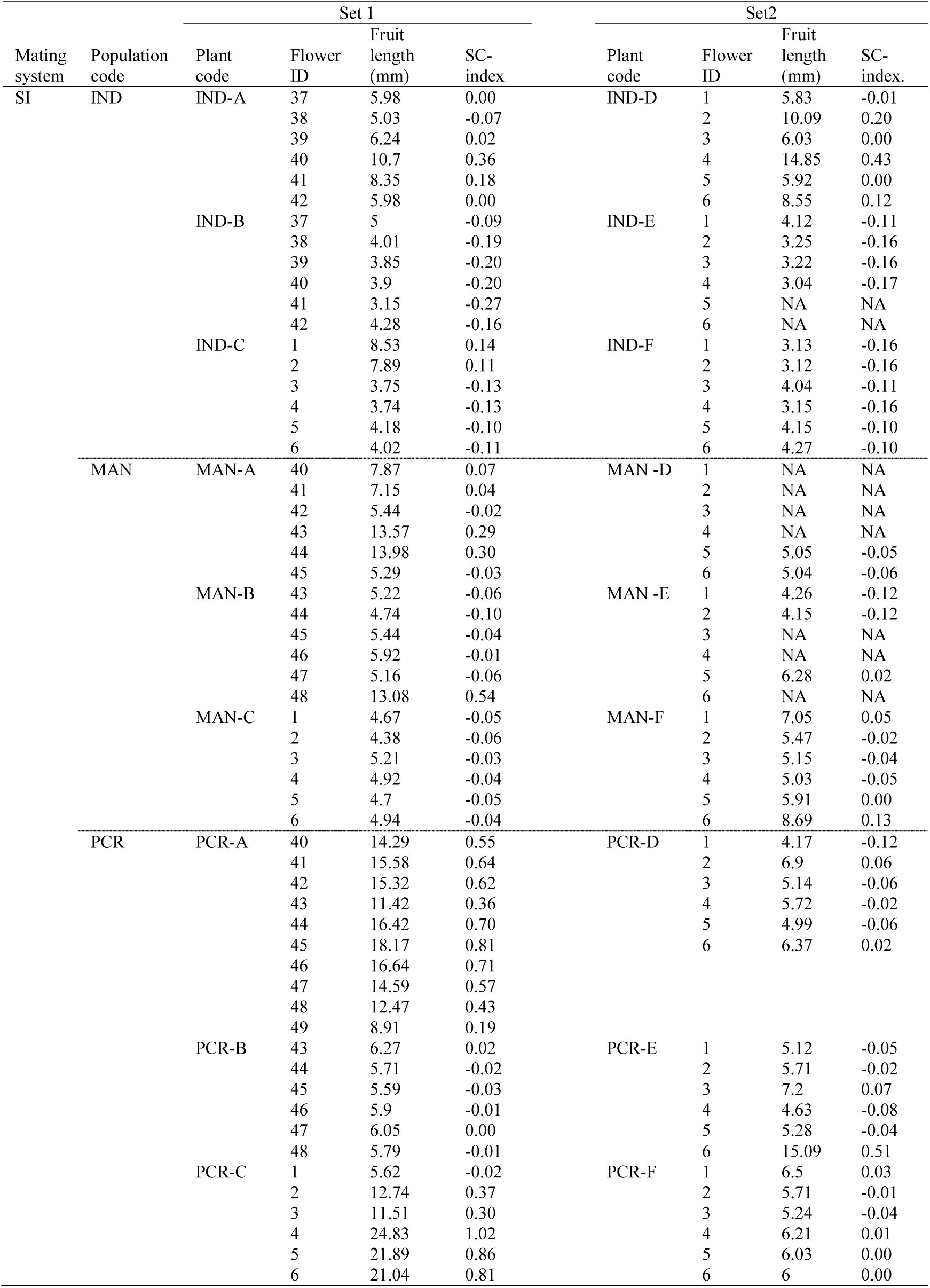
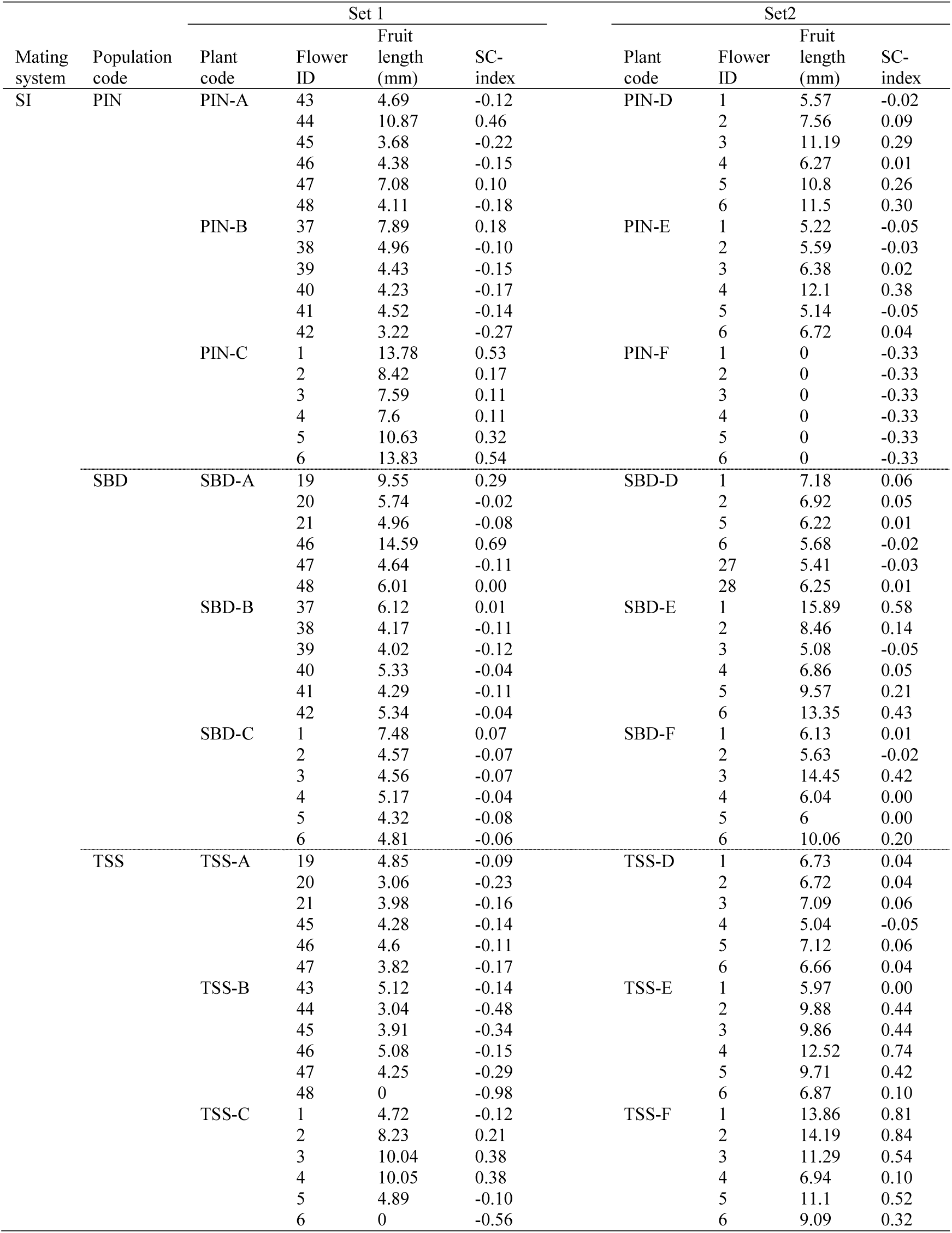
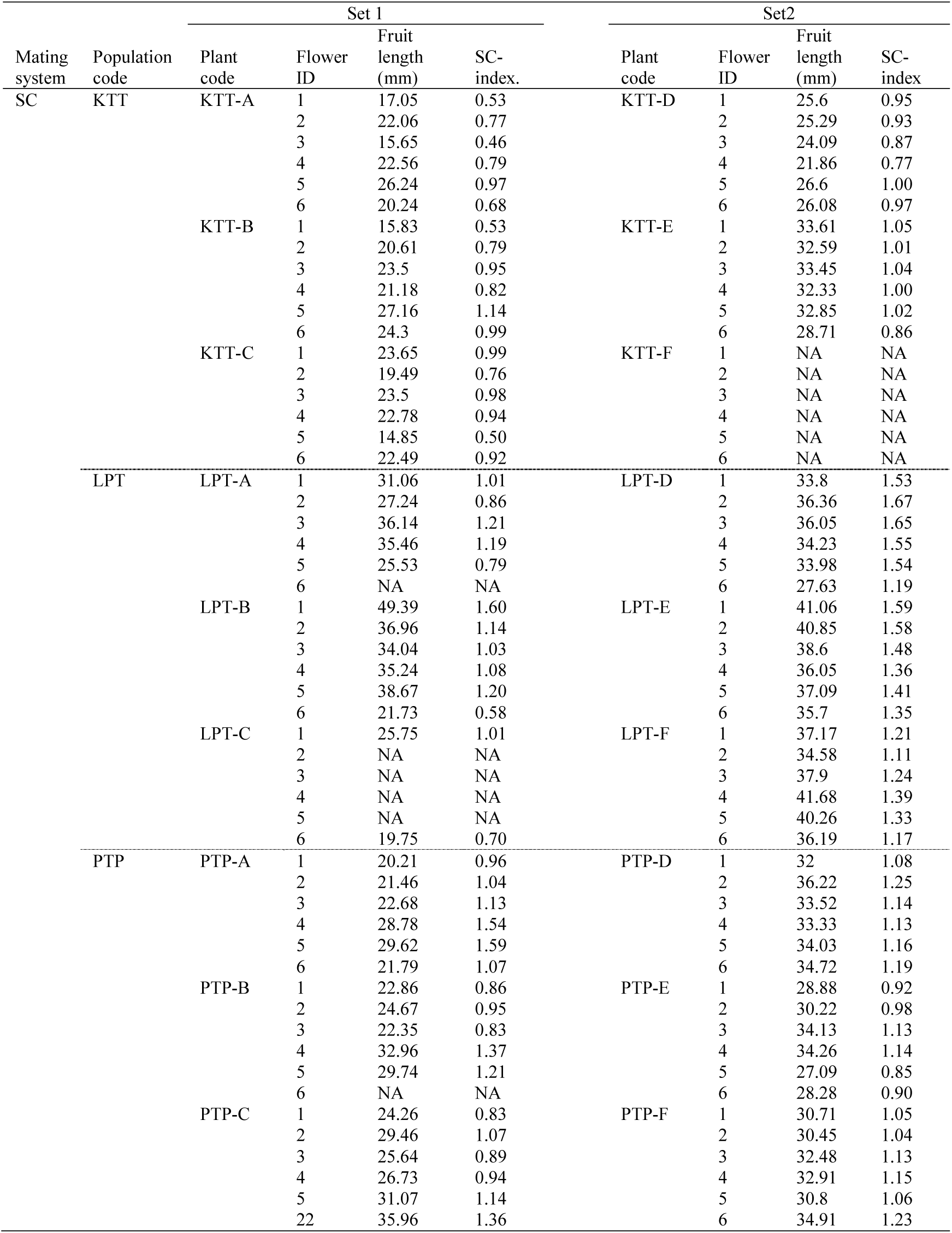
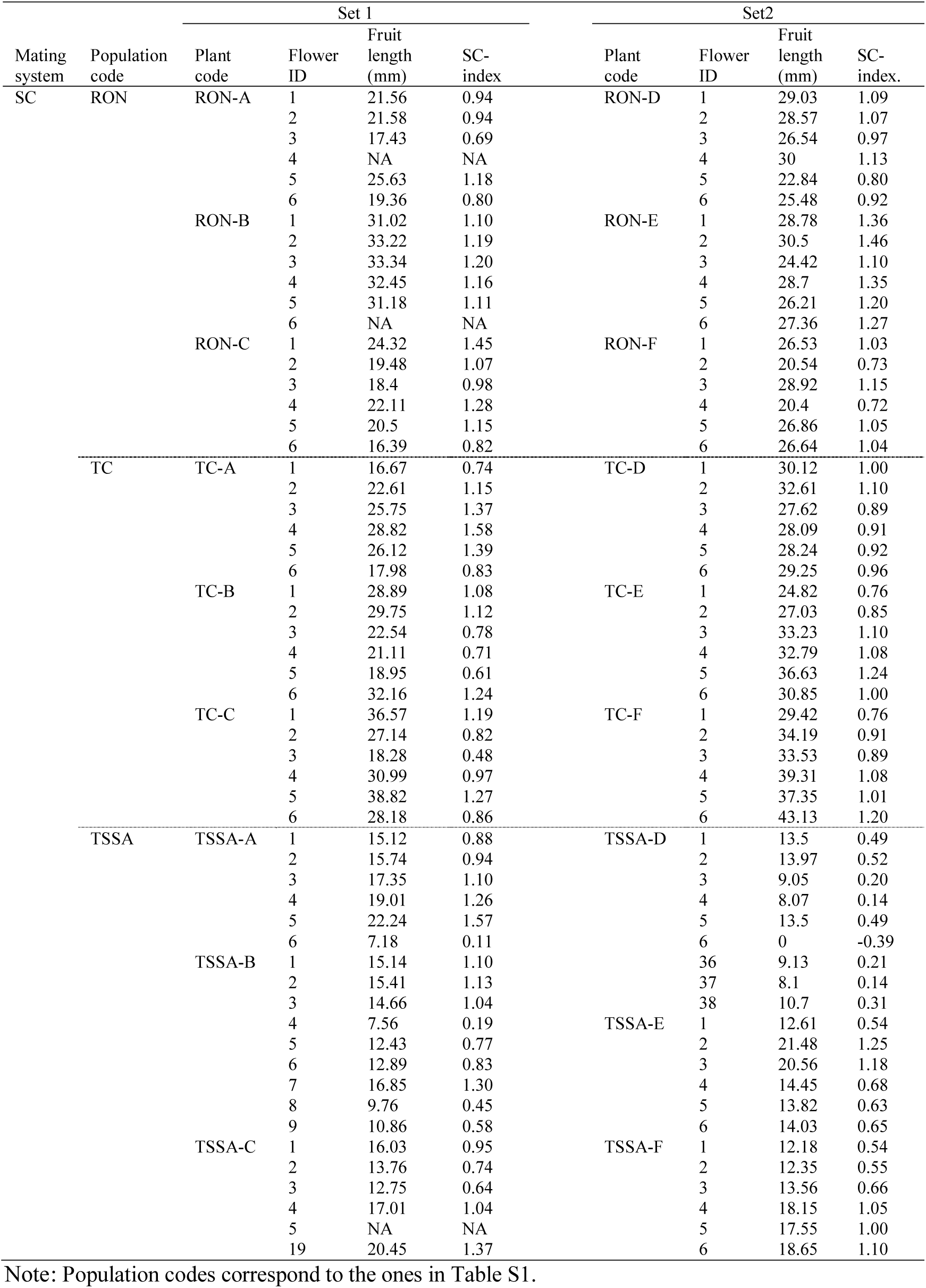
Fruit lengths and corresponding SC-index values after self-pollination of the 72 parental plants in our crossing design (Fig. 1).

**Table S3.** Timeline and progeny numbers for eight batches of cross-progeny.

| Source | Batch number |  |  |  |  |  |  |  | Totals |
| --- | --- | --- | --- | --- | --- | --- | --- | --- | --- |
|  | 1 | 2 | 3 | 4 | 5 | <sup>a</sup> 6 | <sup>b</sup> 7 | <sup>a</sup> 8 |  |
| **Time schedule** |  |  |  |  |  |  |  |  |  |
| Sowing month | 7.2015 | 10.2015 | 9.2016 | 2.2017 | 8.2017 | 8.2017 | 1.2018 | 2.2018 | - |
| First pollination | 9.2015 | 1.2016 | 11.2016 | 4.2017 | 10.2017 | 10.2017 | 4.2018 | 4.2018 | - |
| Last pollination | 3.2016 | 7.2016 | 4.2017 | 9.2017 | 4.2018 | 4.2018 | 8.2018 | 9.2018 | - |
| **Seed and germination** |  |  |  |  |  |  |  |  |  |
| Parental set | A, B, C | A, B, C | D, E, F | D, E, F | A, B, C | A, B, C, D, E, F | A, B, C | A, B, C | - |
| Number of families with at least one seed | 446 | 446 | 452 | 452 | 430 | 211 | 269 | 204 | 2,437 |
| Number of germinated seeds | 419 | 427 | 204 | 206 | 326 | 155 | 164 | 138 | 2,039 |
| **Self-Pollination** |  |  |  |  |  |  |  |  |  |
| Number of plants that could be self-pollinated | 276 | 196 | 202 | 206 | 319 | 145 | 153 | 110 | 1,607 |
| Total number of self-pollinations (in principle, 10 self-pollinations per plant) | 2,564 | 1,855 | 2,045 | 2,137 | 3,247 | 1,503 | 1,483 | 1,146 | 15,980 |
| **Sterility testing** |  |  |  |  |  |  |  |  |  |
| Number of tested plants | 116 | 62 | 95 | 85 | - | - | - | - | 358 |
| Fertile | 114 | 55 | 87 | 75 | - | - | - | - | 331<br>(93.0%) |
| Completely sterile | 0 | 4 | 3 | 3 | - | - | - | - | 10<br>(2.8%) |
| Only female sterile | 0 | 0 | 5 | 7 | - | - | - | - | 12<br>(3.4%) |
| Only male sterile | 2 | 3 | 0 | 0 | - | - | - | - | 5<br>(1.4%) |

Notes: In each batch, we sowed one seed per seed-family
<sup>a</sup> This batch only included progeny from between-breeding-system crosses.
<sup>b</sup> This batch included progeny from crosses between-breeding-system and within-population, but not between populations with the same breeding system.

**Table S4.** Linear mixed model analysis to test the effect of maternal and paternal fruit length and their interaction on progeny fruit length after self-pollination in selfing populations.

| Source | Progeny fruit length after selfing<br>(Gaussian) |  |  |  |  |
| --- | --- | --- | --- | --- | --- |
| **Fixed effects** | df | Estimate | SE | $\chi^2$ | P |
| Maternal fruit length<br>after selfing (MFL) | 1 | 0.32 | 0.22 | **4.97** | **0.026** |
| Paternal fruit length<br>after selfing (PFL) | 1 | 0.32 | 0.22 | **8.50** | **0.004** |
| MFL:PFL | 1 | -0.00 | 0.01 | 0.10 | 0.756 |
| **Random effects** |  |  |  | SD |  |
| Maternal population | 1 |  |  | 1.59 |  |
| Paternal population | 1 |  |  | 1.02 |  |
| Maternal plant<br>(nested in Maternal<br>population) | 1 |  |  | 1.70 |  |
| Paternal plant<br>(nested in Paternal<br>population) | 1 |  |  | 1.40 |  |
| Batch code | 1 |  |  | 0.84 |  |
| Residual | 219 |  |  | 4.22 |  |

Notes: We tested the significance of the fixed effects by comparing models with and without the tested term using likelihood-ratio tests. Significant effects are indicated in bold. We individually calculated fruit length of progeny plants and parental plants based on the median fruit length of multiple fruits per plant prior to analysis.

**Table S5.**
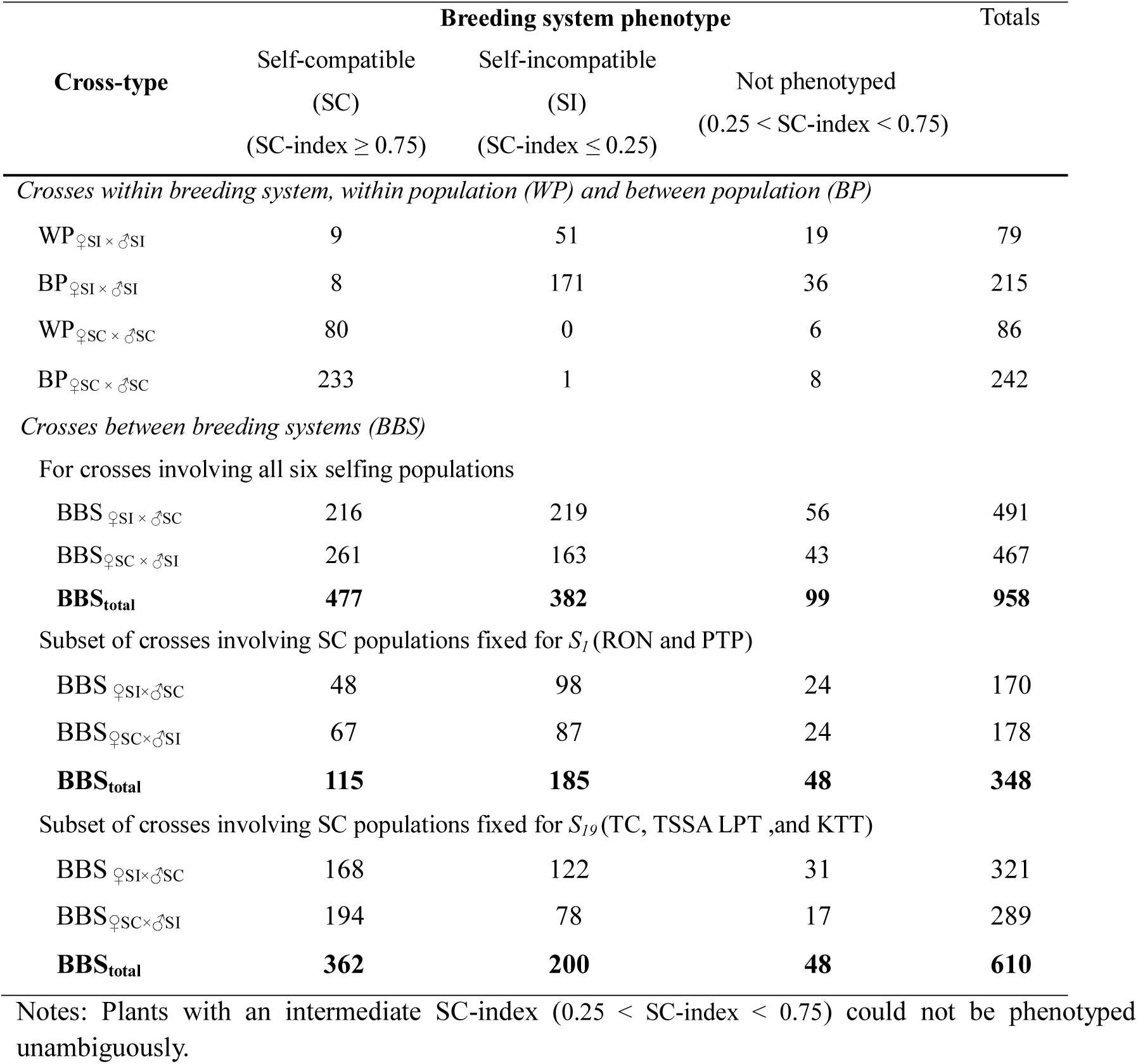
Breeding system phenotype frequencies based on SC-index values for cross-progeny for each cross-type.

**Fig. S1.**
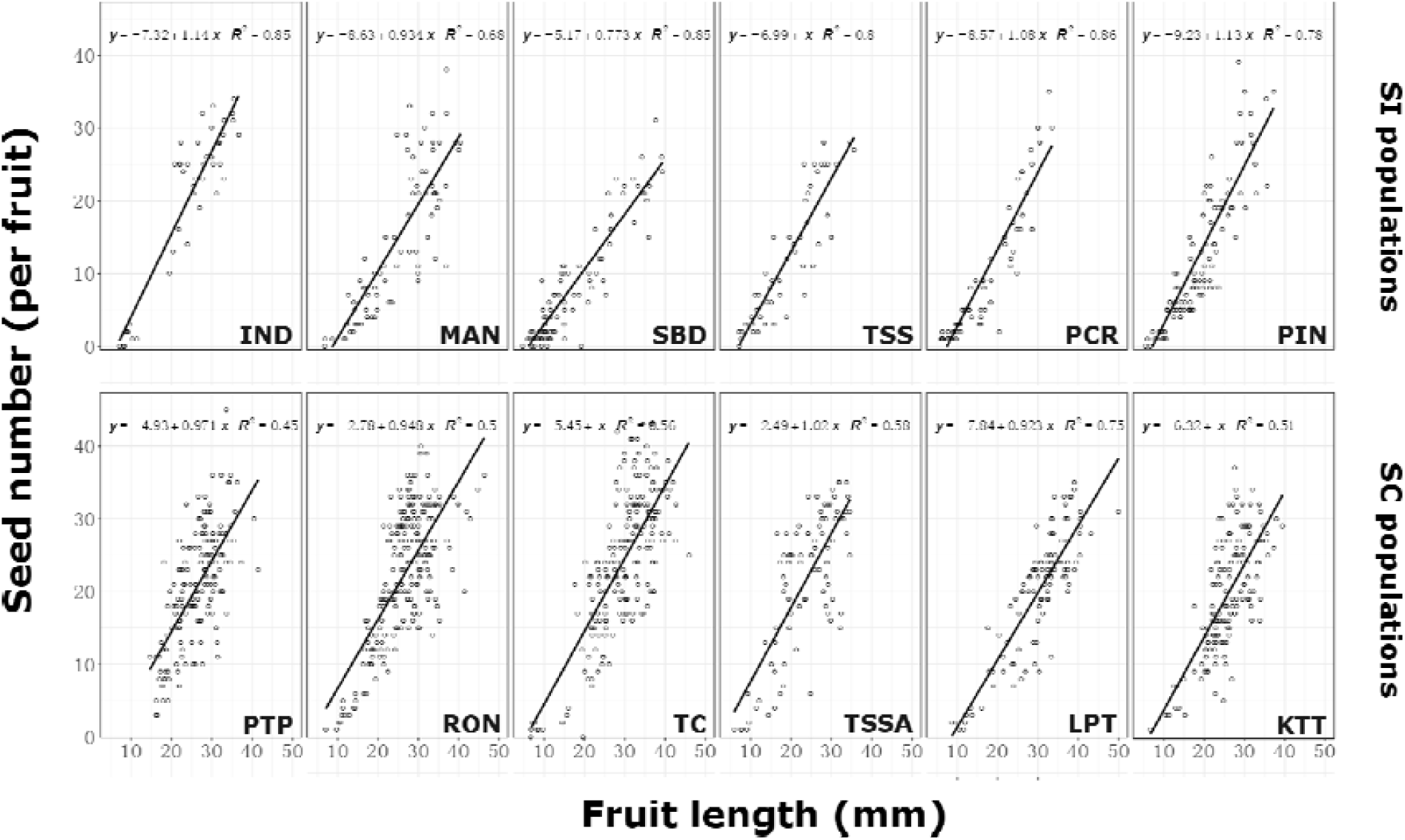
Relationship between fruit length (mm) and seed number per fruit by maternal population. For the subset of the fourth and fifth two batches, we counted seeds per fruit for self-pollinated fruits, to test whether fruit length was a good predictor of seed set. Population codes correspond to those in Table S1). Best-fit regression lines are shown along with their equation and proportion of explained variance R^2^.

**Fig. S2.**
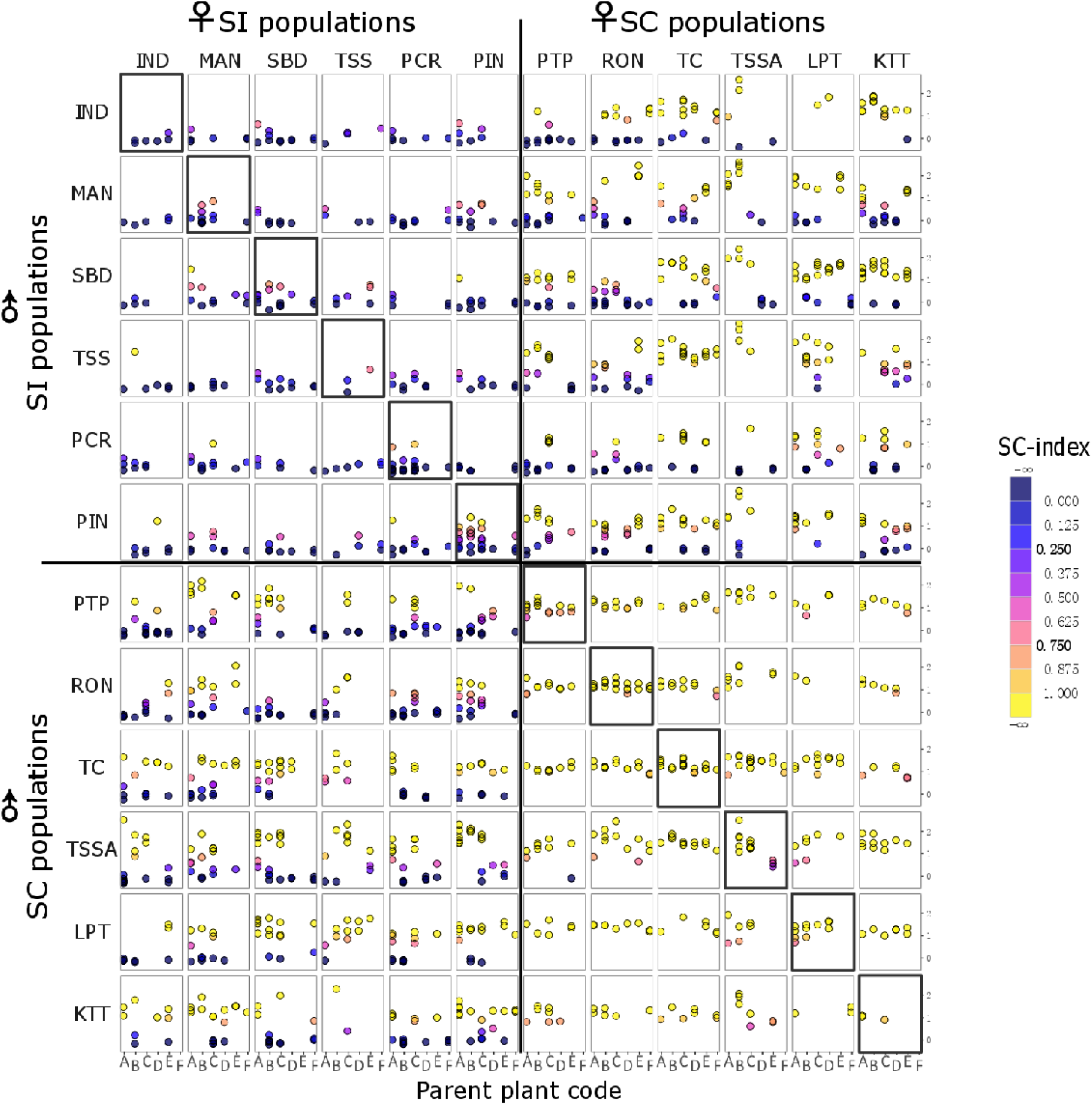
SC-index values for individual cross-progeny by seed family. Crosses between populations were always done between parents with the same plant code (i.e. plants coded A were always crossed with another plant coded A, plants coded B with other plants coded B, et cetera, cf. crossing design in Fig. 1). Population coding corresponds to the coding in Table S1. Diagonal cells marked with an extra outline represent progeny from within-population crosses which resulted from crosses among plants coded A, B and C and among plants coded D, E and F.

**Fig. S3.**
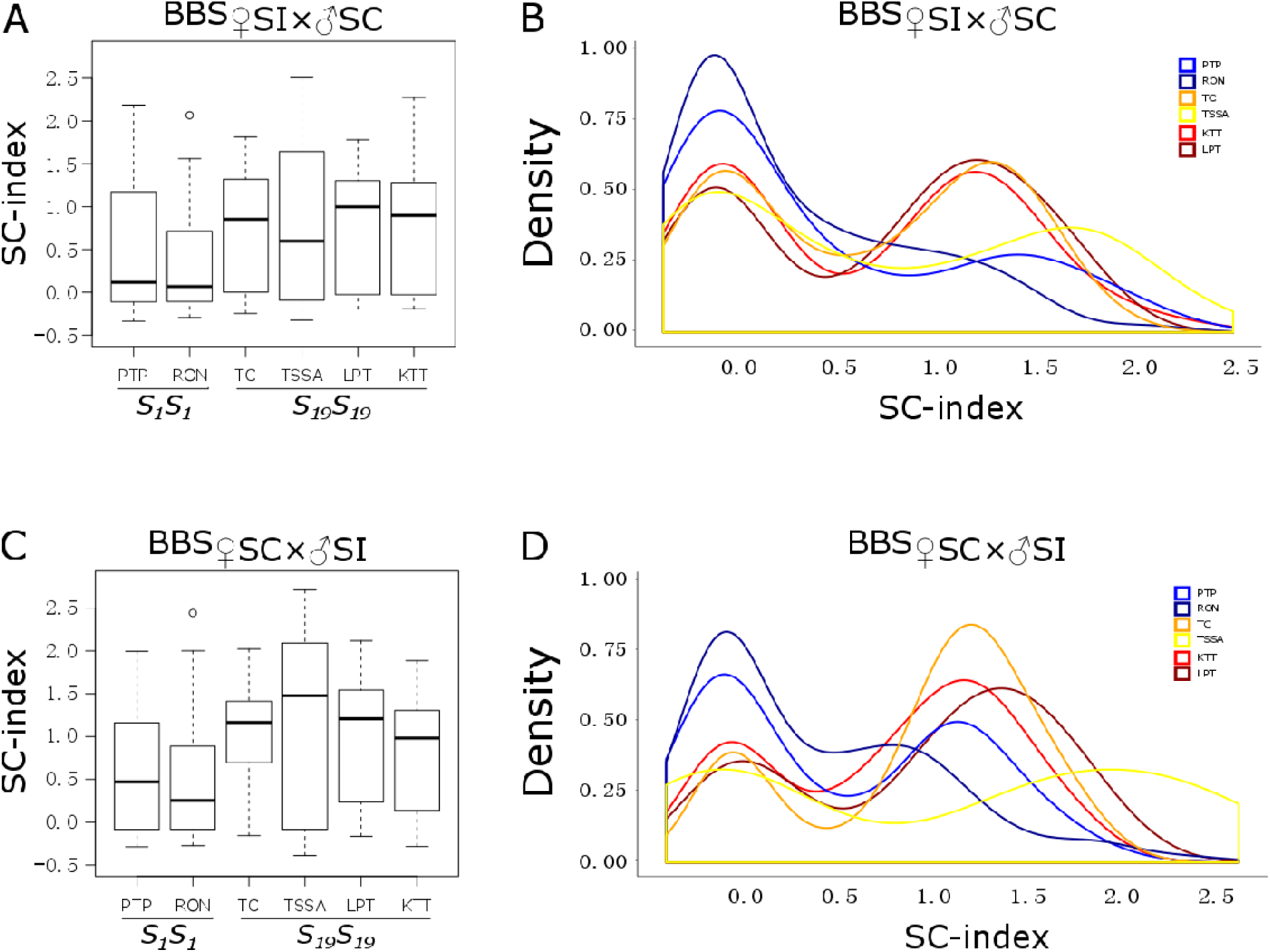
SC-index values for progeny from crosses between breeding systems for the six SC populations grouped by *S*-haplotype. Box plots (A, C) show the interquartile range (IQR) around the median (black horizontal line), with whiskers extending to 1.5×IQR or to the lowest/highest data point within 1.5×IQR. Points beyond whiskers’ limits represent outliers. Density plots (B, D). The *S-*haplotypes of SC parents from PTP and RON are *S_1_S_1_* homozygotes, and from the other four populations *S_19_S_19_*.

